# Progression of recent *Mycobacterium tuberculosis exposure* to active tuberculosis is a highly heritable complex trait driven by 3q23 in Peruvians

**DOI:** 10.1101/401984

**Authors:** Yang Luo, Sara Suliman, Samira Asgari, Tiffany Amariuta, Roger Calderon, Leonid Lecca, Segundo R. León, Judith Jimenez, Rosa Yataco, Carmen Contreras, Jerome T. Galea, Mercedes Becerra, Sergey Nejentsev, Marta Martínez-Bonet, Peter A. Nigrovic, D. Branch Moody, Megan B Murray, Soumya Raychaudhuri

## Abstract

Among 1.8 billion people worldwide infected with *Mycobacterium tuberculosis*, 5-15% are expected to develop active tuberculosis (TB). Approximately half of these will progress to active TB within the first 18 months after infection, presumably because they fail to mount the initial immune response that contains the local bacterial spread. The other half will reactivate their latent infection later in life, likely triggered by a loss of immune competence due to factors such as HIV-associated immunosuppression or ageing. This natural history suggests that undiscovered host genetic factors may control early progression to active TB. Here, we report results from a large genome-wide genetic study of early TB progression. We genotyped a total of 4,002 active TB cases and their household contacts in Peru and quantified genetic heritability 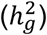 of early TB progression to be 21.2% under the liability scale. Compared to the reported 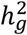 of genome-wide TB susceptibility (15.5%), this result indicates early TB progression has a stronger genetic basis than population-wide TB susceptibility. We identified a novel association between early TB progression and variants located in an enhancer region on chromosome 3q23 (rs73226617, OR=1.19; *P* < 5×10^−8^). We used *in silico* and *in vitro* analyses to identify likely functional variants and target genes, highlighting new candidate mechanisms of host response in early TB progression.

---

The infectious pathogen *Mycobacterium tuberculosis (M.tb)* infects about one quarter of the world’s population^1^. Approximately 5-15% of infected individuals progress to active TB while the vast majority remain infected with viable latent *M.tb* (**Figure 1a**). From the approximately 10.4 million patients with active TB, an estimated ∼1.3 million people died in 2016^2^. Active TB can develop immediately (within the first 18 months) after recent *M.tb* infection or after many years of latency, presumably caused via distinct disease mechanisms. Late progression or TB reactivation is more likely the consequence of acquired immune compromise due to other diseases or ageing, whereas early progression is presumably due to failure in mounting the initial immune response that contains the bacterial spread. Previous studies have indicated a strong heritable component of population-wide TB susceptibility, that includes early disease progression, reactivation and infection^3–5^. But whether early progression has a different genetic architecture compared to population-wide susceptibility has yet to be defined.

**Figure 1.**
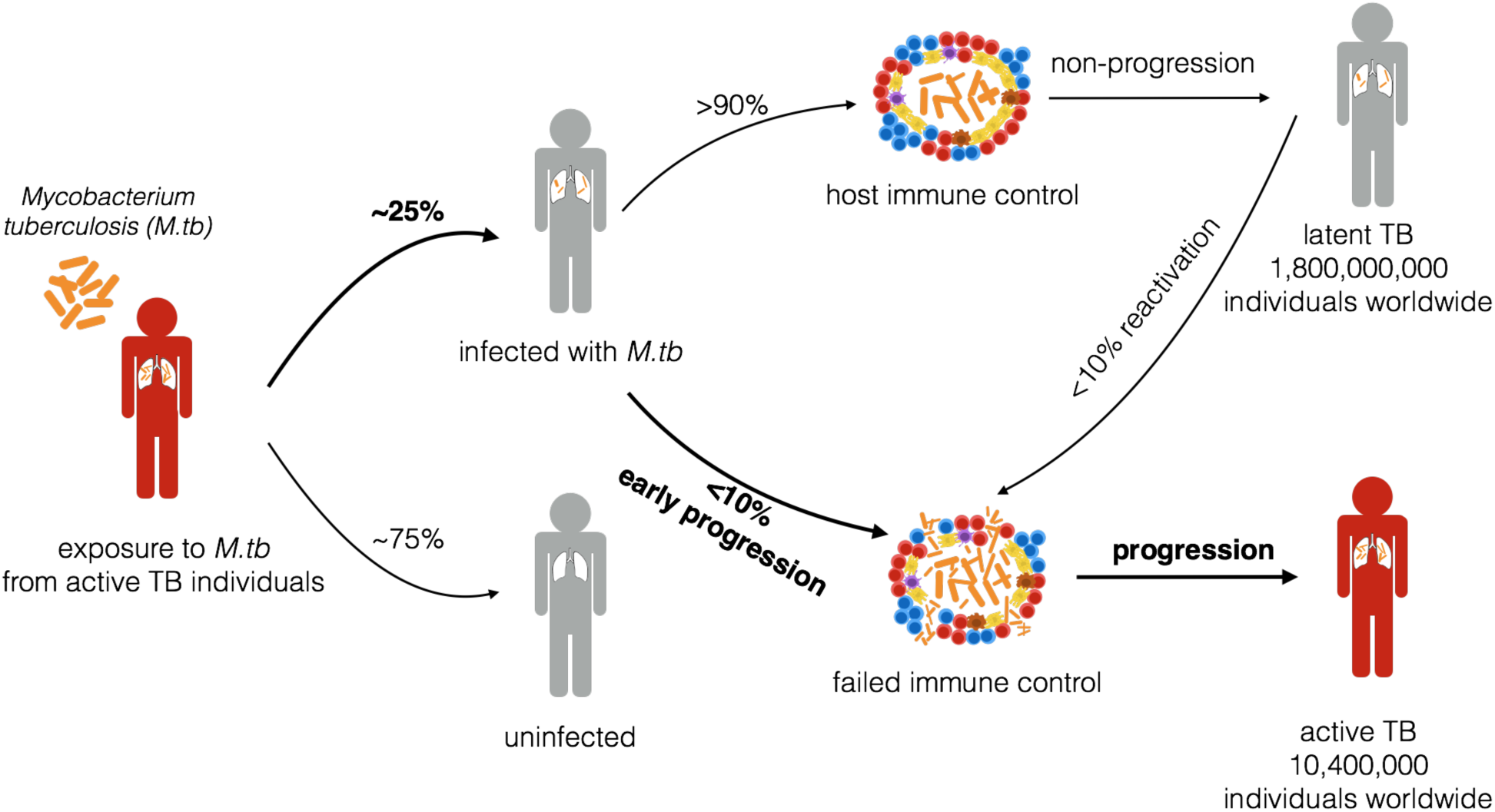
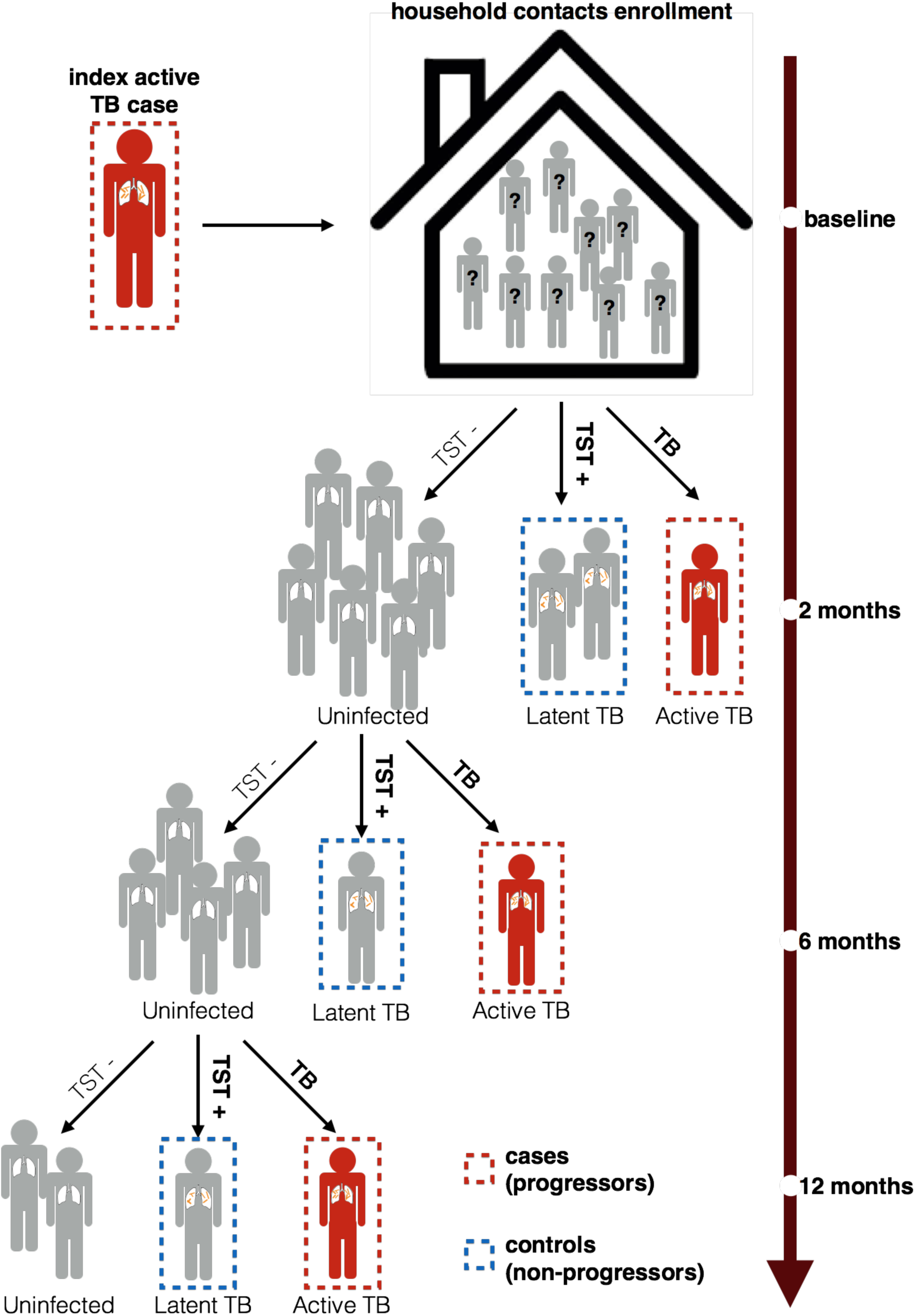
Overview of phases of *Mycobacterium tuberculosis* (*M.tb*) infection and sample collection. (a) Pathophysiology of TB. Major steps following from the initial exposure to *M.tb* are outlined, with the percentages of individuals progressing between steps taken from the WHO TB report^2^. (b) Schema of cohort collection. In this study, we focus on a genetic study between recently exposed active pulmonary TB cases (progressors) and subjects with tuberculin skin test (TST) positive results, who did not progress to active TB (non-progressors). Index cases had sputum with confirmed TB. Controls were recruited in the same household as index cases, with 12 month follow-up periods to confirm infection status using TST.

Reported associations for TB, and other infectious diseases, has to be considered in the context of TB diagnostic criteria and selected control groups^6,7^. To date, genome-wide association studies (GWAS) of TB have compared mixed pools of TB patients with early progression or reactivation, to population controls, who may not have been exposed to *M.tb* at all^8–12^. Hence, known human genetic loci associations with clinical outcomes might represent risk factors for *M.tb* infection, progression from recent *M.tb* exposure to active TB, or reactivation of TB after a period of latency. Infection, progression and reactivation represent pathophysiologically distinct disease transitions likely involving distinct mechanisms of transmission, early innate immune response and control by adaptive immunity. Thus, the study of mixed TB populations using controls of unknown exposure status may underestimate or miss genetic associations for these separate stages of disease.

To identify host factors that drive pulmonary early TB progression, we conducted a large, longitudinal genetic study in Lima, Peru (**Fig. 1b**), where the TB incidence rate is one of the highest in the region^2^. We enrolled patients with microbiologically confirmed pulmonary TB. Within two weeks of enrolling an index patient, we identified their household contacts (HHCs) and screened for infection as measured by a tuberculin skin test (TST) and for signs and symptoms of pulmonary and extra-pulmonary TB. HHCs were re-evaluated at two, six and twelve months. We considered individuals to be early progressors if they are (1) index patients whose *M.tb* isolates shared a molecular fingerprint with isolates from other enrolled patients; (2) HHCs who developed TB disease within one year after exposure to an index patient and (3) index patients who were 40 years old or younger at time of diagnosis. We considered HHCs who were TST positive at baseline or any time during the 12 month follow up period, but who had no previous history of TB disease and remained disease free, as non-progressing controls (**Methods, Figure 1b**). In total, we genotyped 2,175 recently exposed pulmonary TB cases (early progressors) versus 1,827 HHCs with latent tuberculosis infection, who had not progressed to active TB during one year of follow-up (non-progressors), as controls (**Methods, Supplementary Table 1**).

To our knowledge, this represents the most extensive genetic study conducted in Peru to date. Peru is a country with a complex demographic history and underexplored genomic variation. When Spanish conquistadors arrived in the region in the 16th century, Peru was the center of the vast Inca Empire and was inhabited by a large Native American population^13,14^. During the colonial period, Europeans and Africans (brought in as slaves) arrived in large numbers to Peru. After Peru gained its independence in 1821, there was a flow of immigrants from southern China to all regions of Peru as a replacement for slaves^15,16^. As a result, the genetic background of the current Peruvian population is shaped by different levels of admixture between Native Americans, Europeans, African and Asian immigrants that arrived in waves with specific and dated historical antecedents. When compared to individuals from other South American countries^17,18^, Peruvians tend to share a greater genetic similarity with Andean indigenous people such as Quechua and Aymara (**Figure 2, Supplementary Figure 1, Methods**).

**Figure 2.**
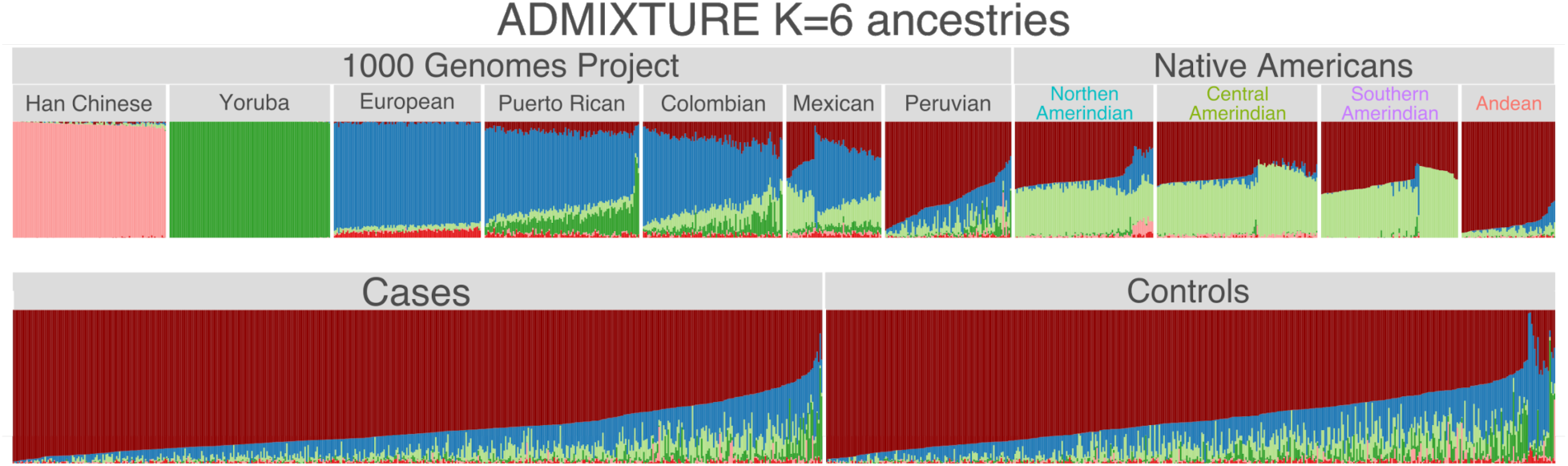
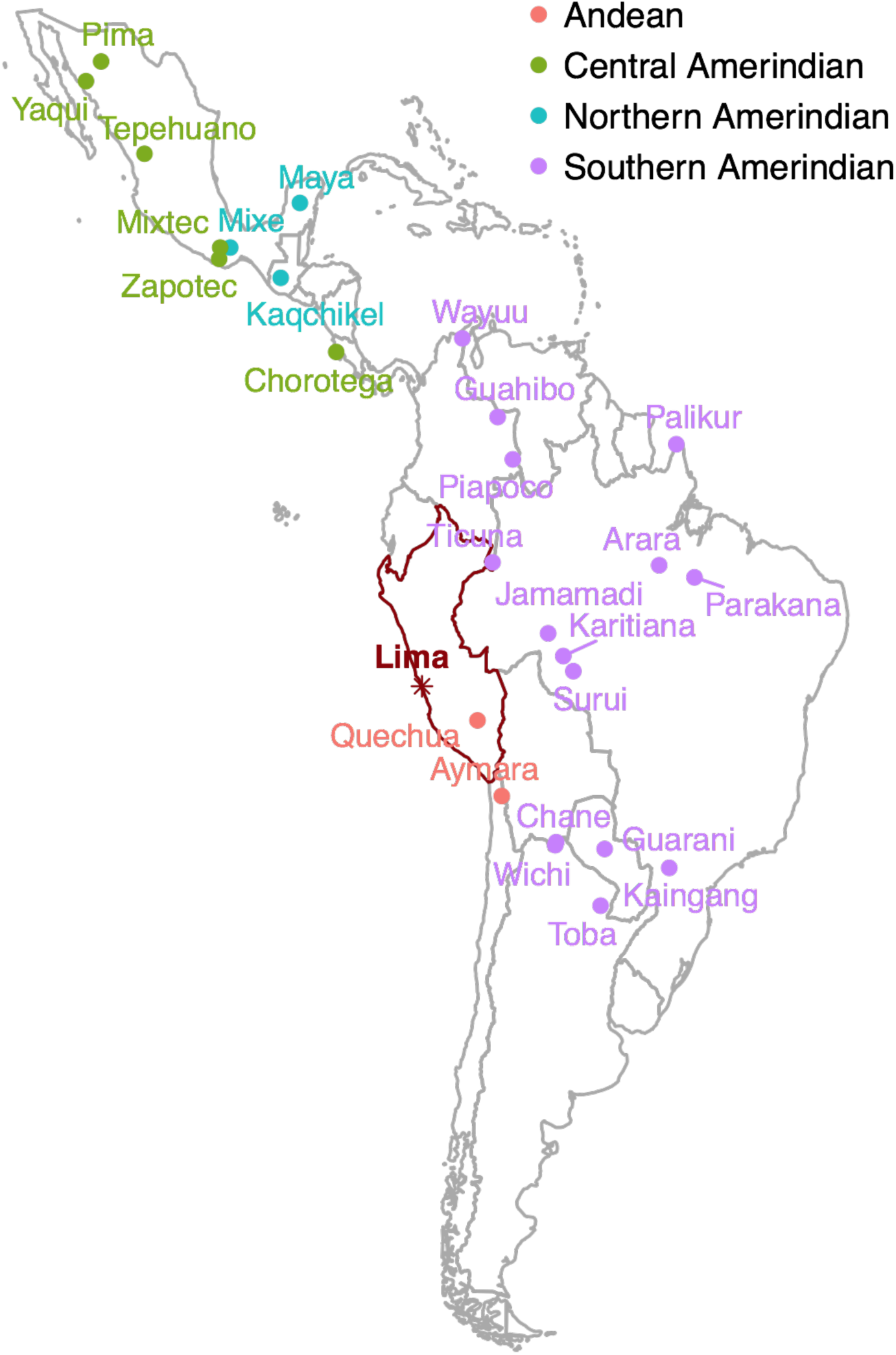
Global ancestry analysis of Peruvian populations. (a) ADMIXTURE plot of admixed individuals and continental reference panels. Each individual is represented as a thin vertical bar. The colors can be interpreted as different ancestries. Reference panels are either from the 1000 Genomes project^17^ (1000G) or Native American individuals collected from *Reich et al. 2012 Nature*^*18*^. Han Chinese are from Beijing, China; Yoruba are from Ibadan, Nigeria; European individuals are Utah Residents (CEPH) with Northern and Western European Ancestry; Puerto Rican samples are from Puerto Rico; Colombian samples are from Medellin, Colombia; Mexican individuals are from Los Angeles, California; Peruvian samples are from Lima, Peru. Northern Amerindian includes individuals from Maya, Mixe and Kaqchikel. Central Amerindian includes individuals from Pima, Zapotec, Mixtec, Yaqui, Chorotega, Tepehuano. Southern Amerindian includes individuals from Piapoc, Karitiana, Surui, Wayuu, Jamaadi, Parakana, Guarani, Kaingang, Ticuna, Palikur, Toba, Arara, Wichi, Chane and Guahibo. Andean population includes Quechua and Aymara. *K* = 6 models are shown above, *K* = 3 through *K* = 15 models are available in **Supplementary Figure 1.** (b) Map of locations of sampled Native American populations^18^.

This unique genetic heritage provides both a challenge and an opportunity for biomedical research. To optimally capture genetic variation, and particularly rare variations in Peruvians, we designed a 712,000-SNP customized array (LIMAArray) with genome-wide coverage based on whole-exome sequencing data from 116 active TB cases (**Methods, Supplementary Table 2, Supplementary Figure 2**). When compared to other more comprehensive genotyping platforms available at the time, LIMAArray showed an approximately 5% increase in imputation accuracy, particularly for population-specific and low-frequency variants (**Supplementary Table 3**). We derived estimated genotypes for ∼8 million variants using the 1000 Genomes Project Phase 3^17^ as the reference panel and tested single marker and rare-variant burden associations with linear mixed models that account for both population stratification and relatedness in the cohort (**Supplementary Figure 3-4**, **Methods**). Genome-wide association results of 2,160 cases and 1,820 controls after quality control (**Methods**) are summarized in **Supplementary Figure 5**. We observed no inflation of test statistics (*λ*_*GC*_= 1.03, *λ* _*GC*_ = 1.00 for common and rare association analyses respectively), which suggests potental biases were strictly controlled in our study.

To investigate the genetic basis of early TB progression, we first estimated its variant-based heritability 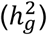. Using GCTA^19^ we estimated 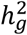 of TB progression to be 21.2% (standard error (s.e.)=0.08, *P* = *2*.64×10^−3^) on the liability scale with assumed incidence rate of 0.05 in the cohort (**Methods**). To avoid biases introduced from calculating genetic relatedness matrices (GRMs) in admixed individuals, we calculated two different GRMs based on admixture-aware relatedness estimation methods^20,21^ and removed related individuals. Both admixture-aware methods reported similar 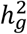 estimates (**Supplementary Table 4**), indicating our reported heritability estimation is robust under different model assumptions. We quantified 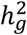 of TB progression and observed a surprisingly strong genetic basis. This degree of heritability is comparable to traits with a well-established genetic basis (**Supplementary Table 5**). For example, GWAS have identified ∼200 risk loci for Crohn’s disease^22,23^, which has a reported 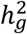 of 28.4% (s.e.=0.02, *P* = 8.*62*×10^−71^)^22^. In contrast, using LD Score regression^24^ on summary statistics from a GWAS of population-wide TB susceptibility in Russia^10^, we estimated the 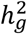 of population-wide TB susceptibility to be 15.5% (s.e.=0.04, *P* = 5.33×10^−5^) with assumed prevalence of 0.04^25^. These data suggest that recently exposed TB progression may have a stronger host genetic basis compared to population-wide TB susceptibility, and larger progression studies may be well-powered to discover additional variants.

We next identified a novel risk locus associated with TB progression on chromosome 3q23, which is comprised of 11 variants in non-coding regions downstream of *RASA2* and upstream of *RNF7* (*P* < 1×10^−5^) (**Figure 3, Supplementary Table 6**). The strongest association was at a genotyped variant rs73226617 (OR=1.19; *P* = 3.93×10^−8^). To test for artifacts and to identify stronger associations that might have been missed due to genotyping and imputation, we first checked the genotype intensity cluster plot of the top associated variant which showed clear separation between genotypes AA, AG and GG (**Supplementary Figure 6**). We then designed individual TaqMan genotyping assays for four top associated variants (**Methods**, **Supplementary Table 7**). We genotyped these four SNPs in 4,002 initial subjects and concluded that all four variants show a high concordance rate (>99%) with imputed genotypes (**Supplementary Table 6**). Because all 11 variants in the risk locus are in high linkage disequilibrium (LD) with each other (**Supplementary Figure 7**), the other imputed variants are also likely to have high imputation quality.

**Figure 3.**
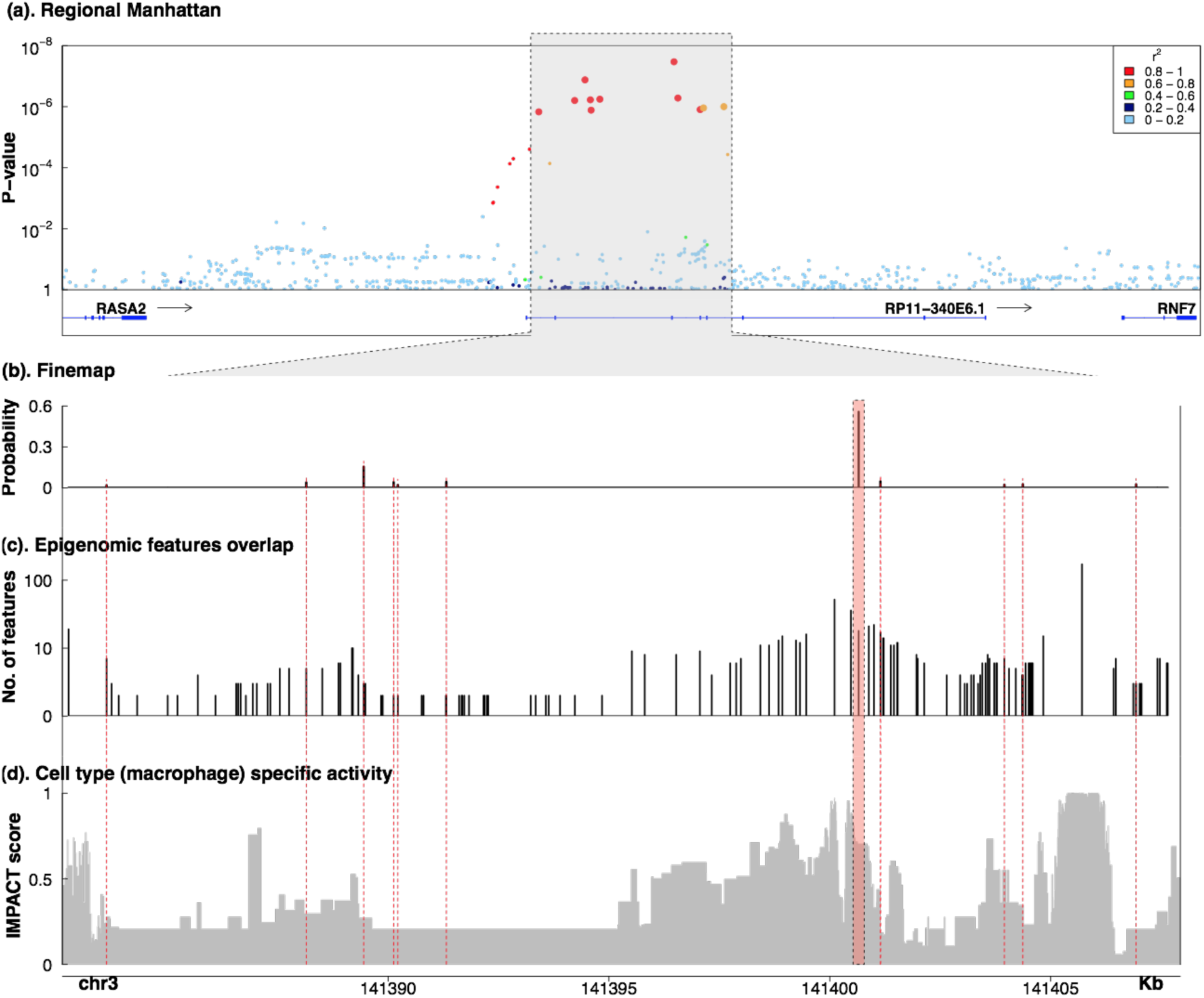
Genome-wide association details of the 3q23 locus. (a) A regional association plot of the 3q23 locus including all genotyped and imputed variants. (b) Fine-mapping posterior probability of all variants in the chr3:140221602-145217859 region. (c) Number of overlaps between all variants in the risk locus and ∼400 epigenetic features. (d) Predicted posterior probability of cell-type specific gene regulatory activity using Inference and Modeling of Phenotype-related ACtive Transcription (IMPACT) based on the epigenetic chromatin signature of binding sites of the transcription factor IRF1. Dashed lines highlights 11 top associated variants. Genotyped variant rs73226617 is highlighted in red.

To determine whether the reported association is specific to TB progression from recent *M.tb* infection or instead derived from reactivation of latent TB, we conducted a case-only analysis removing age from our case selection criteria. This approach is based on the premise that TB cases that share a DNA fingerprint for *M.tb* and HHCs who developed active TB are epidemiologically related while cases in which *M.tb* fingerprints are different might have resulted from remote infection that reactivated during the study assessment^26^. 1,472 out of 2,175 presumed early progressors shared molecular fingerprint of *M.tb isolates* with another case or developed active TB during the one year of follow-up (**Supplementary Figure 8**). Other cases did not have a shared the molecular fingerprint among *M.tb* isolates or did not come from the same household as the index case, leading to a lower degree of certainty in the early progression status of these cases. In this case-only analysis, the top associated signal rs73226617 was nominally associated with early progression (*P* = 0.016, OR=1.09). A heritability analysis restricted to those that shared the same molecular fingerprint or from the same household estimated in a larger 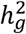 (22.1%, s.e.=0.06, *P* = 1.32×10^−4^) despite the smaller number of samples. These results provide further evidence that the signals we reported at 3q23 are only associated with TB progression after recent exposure to *M.tb* and not from reactivation of latent infection.

We examined our 11 most associated variants for early TB progression identified in the Peruvian cohort in previously published GWAS datasets^8–10,27^ (**Supplementary Table 8**). These SNPs were less frequent (<1%) in the African populations than in the European and Peruvian populations, resulting in lower statistical power to detect association. We therefore examined the SNPs in two previously published Russian^10^ (5,530 TB cases and 5,607 controls) and Icelandic^27^ (4,049 TB cases and 6,543 TST+ controls) GWAS datasets. We observed that the effects in the Russian cohort were similar, as they shared comparable ORs of 1.10 (Peru) and 1.18 (Russia) for rs73226617 (P_Russia_=0.065). In contrast, there was no signal observed in the Icelandic cohort (OR=1.06, P_Iceland_=0.437). Consistent with our previous case-only analysis, the weaker signals observed in both European cohorts indicate that 3q23 is specifically associated with early TB progression. The association signals were therefore most likely diluted due to the inclusion of reactivation cases and non-infected controls in the cohort collection.

We next examined how previously published TB GWAS risk loci are associated in this study. We detected evidence of association in a previously reported TB locus at rs9272785 in the HLA region^27^ (OR=1.04, *P* = 4.49×10^−3^), but did not detect signals at other reported risk loci (**Supplementary Table 9**). Thus, previously reported loci may relate to infection or reactivation phenotypes, rather than early TB progression whereas HLA association may affect both early progression and reactivation. The strongest association observed in the HLA region after imputation (**Method, Supplementary Figure 9**) was rs7739434 located upstream of HLA-A (OR=1.10 *P* = 4.59×10^−7^), indicating a possible HLA class I association with TB progression.

To try to identify which of the variants in our reported risk locus is likely to be the functional polymorphism affecting the risk of pulmonary TB progression, we employed the FINEMAP^28^ software (**Methods**). The 90% credible set includes seven genomic variants, with rs73226617 having the highest posterior probability (0.54), followed by rs58538713 (0.16) and the indel rs148722713 (0.05) among 713 variants in the region (**Supplementary Table 6**). To identify likely functional variants and target genes, we employed a method called IMPACT (Inference and Modeling of Phenotype-related ACtive Transcription)^29^. Briefly, IMPACT identifies regions predicted to be involved in transcriptional regulatory processes related to a cell-type-specific key transcription factor (**Method, Figure 3**) by leveraging information from nearly 400 *in silico* epigenomic and sequence annotations from public databases (**Supplementary Table 10**). We trained IMPACT on the epigenetic chromatin signature of binding sites of the transcription factor IRF1 to identify active regulatory regions specific to macrophages. Among 11 variants in the risk locus, the leading associated variant rs73226617 had the highest predicted probability (0.704) of lying in an active macrophage-specific regulatory region. Overexpression of *IRF1*, along with other Type I interferon response genes, was detected early in tuberculosis contacts who progressed to active disease^30,31^. Overall, we saw an enrichment of the interferon response factor in the 3q23 locus (**Figure 3d**). We then performed electrophoretic mobility shift assays (EMSA) and luciferase assays to functionally identify the most likely causal variant among the seven variants that constitute 90% credible set (**Method**). EMSA tests whether the variants differentially bound nuclear complexes in an allele-specific manner. Four variants (rs73226617, rs148722713, rs11710569 and rs73226608) showed differential EMSA signals in the risk variants that were suppressed with unlabeled probes, consistent with allele-specific protein binding in the Jurkat76 cell line. We performed luciferase assays on the four candidate variants but failed to detect allele-specific enhancer activity on human embryonic kidney (HEK293T) cells (**Methods**, **Supplementary Figure 10**). This negative result may be driven by the sensitivity limits of the assay or the variants having cell-type-specific activities which might not have been captured by the designed assays. Using GeneHancer^32^ version 4.7, the most plausible target genes include *RNF7* (GH03I141681, gene-enhancer score =11.7, 56,393 base pairs (bp) from the top associated variant rs73226617). RNF7 is a highly conserved ring finger protein. It is an essential subunit of SKP1-cullin/CDC53-F box protein ubiquitin ligases, which are a part of the protein degradation machinery important for cell cycle progression and signal transduction. Among its related pathways are innate immune system and class I MHC mediated antigen processing and presentation. Another candidate target gene located near the risk locus is *RASA2*^*32*^, which lies 66,469 bp upstream from our top associated variant and is expressed in human lung cells^33^. The RASA2 protein is a member of the GAP1 family of GTPase-activating proteins (GAPs) that is involved in cellular proliferation and differentiation. It has been indicated that *RASA2* acts as a regulator of alveolar macrophage activation^34^. Interestingly, previously reported TB-associated gene *ASAP1*^*10*^ also encodes GAPs, indicating that the family of GAPs may play an important role in TB pathogenesis.

Overall, our results argue that rapid TB progression is a highly heritable trait, comparable to other human diseases with an established genetic origin. More generally, these results begin to address general questions about genomic approaches to infectious diseases, which have lagged behind other disease in terms of locus discovery in comparison to other complex traits (**Supplementary Table 5**). Infections, especially chronic infectious diseases, play out in highly distinct phases that involve exposure, crossing epithelial boundaries, pathogen expansion, locating a host niche, and in the case of TB, decades-long persistence, reactivation and re-transmission. Each of these stages can be controlled by distinct host factors. Our analysis indicates that progression from recent *M.tb* exposure to active TB has a different genetic basis compared to TB reactivation. Specific analysis of clinical progression as a distinct phase allows for a more powerful detection of risk factors for an equal number of samples, as compared to case-control studies, which are an amalgamation of different phenotypes. Thus, this work argues strongly that while detailed, stage-specific phenotypic profiling may be more costly, it may offer key advantages for infectious disease genetic studies. Specifically, it allows for precise phenotype definitions, greater heritability and identification of biological targets with specific implications. Therefore detailed phenotypic profiling should become a general strategy for future genetic studies of infectious diseases.

## Methods

### Ethics statement

We recruited 4,002 subjects from a large catchment area of Lima, Peru that included 20 urban districts and approximately 3.3 million residents to donate a blood sample for use in our study. We obtained written informed consent from all the participants. The study protocol was approved by the Institutional Review Board of Harvard School of Public Health and by the Research Ethics Committee of the National Institute of Health of Peru.

### Preparation of genome-wide genetic data

We enrolled index cases as adults (aged 15 and older) who presented with clinically suspected pulmonary TB at any of 106 participating health centers. We excluded patients who resided outside the catchment area, who had received treatment for TB before and those who were unable to give informed consent. Pulmonary TB patients have been diagnosed by the presence of acid fast bacilli in sputum smear or a positive *M.tb* culture at any time from enrollment to the end of treatment. All cultures of the index cases were genotyped using mycobacterial interspersed repetitive units-variable number of tandem repeats (MIRU-VNTR). Within two weeks of enrolling an index patient, we enrolled his or her household contacts (HHCs). The *M.tb* status was determined using the Tuberculin Skin Test (TST). All HHCs were evaluated for signs and symptoms of pulmonary and extra-pulmonary TB disease at two, six and 12 months after enrollment. To select cases who were likely to have recently exposed TB, we chose HIV-negative, culture-positive, drug-sensitive pulmonary TB cases from one of three groups: (1) exposed HHCs who developed active TB during a 12 month follow up period; (2) index patients whose *M.tb* isolates shared a molecular fingerprint with isolates from other enrolled patients and (3) index patients who were 40 years old or younger at time of diagnosis. To maximize the likelihood that controls were exposed to *M.tb* but did not develop active disease, we chose them from among TST positive HHCs with no previous history of TB disease, and who remained disease free at at the time of recruitment both by directly re-contacting individuals to inquire about their latest medical history and by checking their names against lists of notified TB patients at all of the 106 health clinics. Where possible, we chose controls who are less than second-degree related to the index cases.

### Customized Axiom array for Peruvian populations

We developed a custom array (LIMAArray) based on whole-exome sequencing data from 116 active TB cases to optimize the capture of genome-wide genetic variation in Peruvians. Many markers were included because of known associations with, or possible roles in, phenotypic variation, particularly TB-related (**Supplementary Table 11**). The array also includes coding variants across a range of minor allele frequencies (MAFs), including rare markers (<1% MAF), and markers that provide good genome-wide coverage for imputation in Peruvian populations in the common (>5%), low frequency (1-5%) and rare (0.5-1%) MAF ranges (**Supplementary Table 3**). This approach allowed the detection of rare population specific coding variants and those which predisposed individuals to TB risk.

### Genotyping and quality control

We extracted genomic DNA from whole blood of the participating subjects. Genotyping of all samples was performed using our customized Affymetrix LIMAArray. Genotypes were called in a total of 4,002 samples using the apt-genotype-axiom^35^. Individuals were excluded if they were missing more than 5% of the genotype data, had an excess of heterozygous genotypes (±3.5 standard deviations, **Supplementary Table 12**), duplicated with identity-by-state >0.9 or index cases with age at diagnosis greater than 40 years old. After excluding these individuals, we excluded variants with a call rate less than 95%, with duplicated position markers, those with a batch effect (*P* < 1×10^−5^), Hardy-Weinberg (HWE) P-value below 10^−5^ in controls, and a missing rate per SNP difference in cases and controls greater than 10^−5^ (**Supplementary Table 13**). In total, there were 3,980 samples and 677,232 SNPs left for imputation and association analyses after quality control.

### Imputation and association analyses

The genotyped data were pre-phased using SHAPEIT2^36^. IMPUTE2^37^ was then used to impute genotypes at untyped genetic variants using the 1000 Genomes Project Phase 3 dataset^17^ as a reference panel. For chromosome X, males are coded as diploid. That is male genotypes are coded as 0/2 and females genotypes are coded as 0/1/2. HLA imputation was performed using SNP2HLA^38^ and a multi-ethnic HLA imputation reference pane^39^. Imputed SNPs were excluded if the imputation quality score *r*^2^ was less than 0.4, HWE P-value < 10^−5^ in controls or a missing rate per SNP greater than 5%. After filtering, 7,756,401 SNPs were left for further association analyses.

Common single variant associations were tested with a linear mixed model (LMM) implemented in GEMMA^40^ version 0.94.1 on genotype likelihood from imputation assuming an additive genetic model. We use the genetic relatedness matrix (GRM) as random effects to correct for cryptic relatedness between collected individuals. Sex and age were included as fixed effects to correct for population stratification (**Supplementary Figure 2**). The GRM was obtained from an LD-pruned (r^2^< 0.2), with MAF ≥1% after removing large high-LD regions^41^ (**Supplementary Table 14)** dataset of 154,660 SNPs using GEMMA^42^ version 0.7-1.

Gene-based rare variant (MAF<1%) burden test was performed using GMMAT^42^ version 0.7-1, a generalized linear mixed model framework. For each gene *j*, we aggregated the information for multiple rare variants into a single burden score 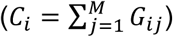 for each subject *i*. Where *G*_*ij*_ denotes the allele counts {0,1,2} for m variants in the gene. The genomic control inflation factor (*λ*_*GC*_) for variants after imputation was 1.03 and 1.00 for common and rare association study respectively (**Supplementary Figure 4**), indicating that we have successfully controlled for any residual population structure or cryptic relatedness between genotyped samples.

To avoid false-positive signals due to population stratification and heterogeneity of effects due to differential LD in admixed populations, we also computed GRMs based on methods^20,21^ that account for inflation of identity-by-state statistics due to admixture LD. LMM with admixture-aware GRMs resulted in numerically similar association statistics to those from unadjusted analyses (**Supplementary Table 15**).

To identify likely causal variants in the identified risk locus, we used FINEMAP^28^ method to calculate marginal likelihoods and Bayes factor for each variant assuming that there is one true causal variant in the region, and it has been included in the analysis and has been well imputed (--n-causal-max 1). We used the in-sample LD scores calculated using LDstore^43^ to further increase the accuracy of the fine-mapping analysis.

### TaqMan SNPs and Genotyping

Selection of SNPs in the 3q23 locus was conducted based on information from the dbSNP database (http://www.ncbi.nlm.nih.gov/projects/SNP/). Two polymorphisms rs73226617, rs73226619, rs73239724 and rs73226608 were included for the genotyping tests. Real-time PCR using the following calculations: 2.5uL Genotyping Master Mix, 0.25uL SNP Assay-probes, and 2.25uL DNA template (at 5ng/uL= 11.25ng total). Thermal cycling conditions were as follows: 60C 30secs Pre-read, 95 °C for 10 min, followed by 40 cycles at 95°C for 15 s and at 60 °C for 1 min, then 60C 30secs Post-read. Genotyping of the polymorphisms was carried out using the 5’ exonuclease TaqMan Allelic Discrimination assay, which was performed utilizing minor groove binder probes fluorescently labeled with VIC or FAM and the protocol recommended by the supplier (Applied Biosystems, Foster City, CA, USA). Analysis for interpretation was performed with Via7 software and Taqman Genotyper software calls. Per variant concordance rate was obtained by comparing genotypes obtained from imputation and from TaqMan assays (**Supplementary Table 6**)

### Heritability estimation

The genetic heritability based on genome-wide markers 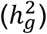 was first estimated from the genetic relatedness matrix (GRM) after removing related individuals (--grm-cutoff 0.125) and corrected for population stratifications using the top 10 principal component (--qcovar), as implemented in GCTA^19,44^. Among a total of 14,044 enrolled HHCs, 692 progressed to active TB. Based on these numbers, we estimated the incidence rate in the Lima cohort for recent TB progression is 5%. Using this rate, we report 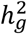 on the liability scale to be 0.21 (s.e. = 0.07). If the true prevalence was in fact half as high, our estimate would instead be 0.17 (s.e. = 0.02); if twice as high, 0.26 (s.e. = 0.09). 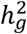 on the observed scale is 0.24 (s.e. = 0.09).

### *In silico* functional annotation of candidate causal variants

We combined multiple sources of *in silico* genome-wide functional annotations from publicly available databases to help identify potential functional variants and target genes in the 3q23 novel risk locus. To investigate functional elements enriched across the region encompassing the strongest candidate causal variants,We aggregated approximately 400 genomic and sequence annotations including cell-type-specific annotation types such as ATAC-seq, DNase-seq, FAIRE-seq, HiChIP-H3K27ac, HiChIP-CTCF, polymerase and elongation factor ChIP-seq, and histone modification ChIP-seq, as well as cell-type-nonspecific annotations such as conservation, coding annotation, and distance to TSS. A list of all included resources is summarized in **Supplementary Table 10**.

The influence of candidate causal variants on transcription factor binding sites was identified using HaploReg^45^ version 4.1. Among possible motif changes, IRF1 is a key transcriptional regulator (TF) that plays critical roles in activation of macrophages by proinflammatory signals such as interferon-γ and highly relevant to tuberculosis pathogenesis^46,47^. We subsequently determined genome-wide TF occupancy from publicly available ChIP-seq of IRF1 (**Supplementary Table 16**), a key regulator of monocyte-derived macrophages (GSE100381)^48^. Briefly, CD14+ human monocytes were purified from PBMCs and then treated with a macrophage colony-stimulating factor (M-CSF). ChIP-seq peaks were called by macs^49^ [v1.4.2 20120305] (FDR<0.05). Using IMPACT (Inference and Modeling of Phenotype-related ACtive Transcription)^29^, we built a model that predicts TF binding on a motif by learning the epigenomic profiles of the TF binding sites. We train IMPACT on gold standard regulatory and non-regulatory elements of IRF1. To build the regulatory class, we scanned the IRF1 ChIP-seq peaks, mentioned above, for matches to the IRF1 binding motif, using HOMER^50^ [v4.8.3] and retained the highest scoring match for each ChIP-seq peak. To build the non-regulatory class, we scanned the entire genome for IRF1 motif matches, again using HOMER, and selected motif matches with no overlap with IRF1 ChIP-seq peaks. IMPACT learns the feature in 10-fold cross validation (CV) of the complete sets of regulatory and non-regulatory elements. We scored regions of interest with the learned from this CV.

### Electrophoretic Mobility Shift Assay (EMSA)

Frozen cell pellets from the Jurkat76 cell line (ATCC) were used for preparation of nuclear extracts using NE-PER Nuclear and Cytoplasmic Extraction reagent (ThermoFisher) according to the manufacturer’s instructions, then dialyzed overnight at 4°C with gentle stirring in 1L of pre-cooled dialysis buffer (10% glycerol, 10mM Tris pH 7.5, 50mM KCl, 200mM NaCl, 1mM di-thiothreitol, 1mM phenylmethanesulfonyl fluoride). Samples were quantified using BCA Protein Assay Kit (ThermoFirsher) and stored in 1X Halt protease inhibitor cocktail (ThermoFisher) at −80°C until use. We designed single stranded oligonucleotides (30-34bp) corresponding to each set of alleles (Integrated DNA Technologies, **Supplementary Table 17**), and biotinylated the forward and reverse sequences separately using the Biotin 3’End DNA Labeling Kit (ThermoFisher Scientific) following the manufacturer’s instructions. Single stranded probes were annealed by incubation for 5 minutes at 95°C followed by 1 hour at room temperature. EMSA reactions were performed using the LightShift Chemiluminscent EMSA kit (ThermoFisher). Binding reactions were performed in a volume of 20μL: 2μL of 10 × binding buffer, 16μg nuclear extract, 2.5% glycerol, 5mM MgCl2, 0.05% Nonidet P-40 and 50ng Poly dI:dC as a non-specific DNA competitor, and 20fmol of biotinylated probes with or without unlabeled competitor probes at 200 fold molar excess. The assay was performed as previously described^51^

### Luciferase reporter assays

We designed double stranded oligonucleotides matching the probes used for EMSA and flanked by either BglII or BamHI restriction sites **(Supplementary Table 18)** for cloning either upstream or downstream of the firefly Luciferase (*Luc*) gene in the pGL3 promoter reporter vector (Promega), respectively. Double stranded inserts were cloned into the pGL3 vector following standard cloning protocols and verified by colony PCR reactions **(Supplementary Table 19)** as well as plasmid Sanger sequencing (GENEWIZ). Plasmids with inserts cloned in-sense with the luciferase promoter sequence were expanded and purified using PureLink HiPure Plasmid Miniprep Kits (Thermofisher Scientific). For transfection, 10^4^ human embryonic kidney (HEK293T) cells were plated per well in 60μL Dulbecco’s Modified Eagle Medium-10 media (DMEM, Gibco, 10% fetal bovine serum, 1x penicillin-streptomycin) in flat-bottom 96-well plates and transfected with 500ng of plasmids (4:1 of pGL3:pRL-TK), using lipofectamine LTX Reagent with PLUS (Thermofisher) according to the manufacturer’s instructions. Transfected cells were incubated for 18-20 hours at 37°C, then analyzed using two-step Dual-Glo® Luciferase Assay System (Promega) and read on Synergy H1 Hybrid Multi-Mode Reader (BioTek). Luminscence is reported as the ratio of firefly (pGL3) to renilla (pRL-TK) luciferase luminescence, normalized to pGL3. We compared the pooled averages of triplicates per plate, paired by transfection plate from 3-10 independent experiments for each variant using a Wilcoxon signed-rank test.

## Acknowledgements

The study was supported by the National Institutes of Health (NIH) TB Research Unit Network, Grant U19 AI111224-01. The content is solely the responsibility of the authors and does not necessarily represent the official views of the NIH. S.N. was supported by MRC (MR/M012328/1), the ERC Starting grant (260477) and the National Institute for Health Research (NIHR) Cambridge Biomedical Research Centre. The authors thank Garðar Sveinbjornsson, Patrick Sulem, Ingileif Jonsdottir and Kari Stefansson at deCODE genetics, Reykjavik, Iceland, for validating the association of rs73226617 with TB progression in the Icelandic population.

## Author contributions

Y.L. designed the genotyping array, performed statistical analysis of the GWAS data and wrote the first draft of the manuscript. S.S. performed the EMSA and luciferase assays experiments. S.A. carried out the rare association studies of the GWAS data. T.A. implemented the IMPACT model. R.C., L.L., S.R. L., J. J., R. Y., C.C., J T. G., M.B. and M.B.M. participated in study design, protocol development and sample collection. S.N. contributed the Russian data for the meta and heritability analysis. M. MB. and P.A.N. helped to develop the protocols for EMSA and luciferase assays experiments. D.B.M supervised the EMSA and luciferase assay experiments. M.B.M. participated in study design, protocol development, and study conception. S.R. conceived and supervised the study. All authors contributed to the writing of the manuscript.

